# A scalable and unbiased discordance metric with *H*_+_

**DOI:** 10.1101/2022.02.03.479015

**Authors:** Nathan Dyjack, Daniel N. Baker, Vladimir Braverman, Ben Langmead, Stephanie C. Hicks

## Abstract

A standard unsupervised analysis is to cluster observations into discrete groups using a dissimilarity measure, such as Euclidean distance. If there does not exist a ground-truth label for each observation necessary for external validity metrics, then internal validity metrics, such as the tightness or consistency of the cluster, are often used. However, the interpretation of these internal metrics can be problematic when using different dissimilarity measures as they have different magnitudes and ranges of values that they span. To address this problem, previous work introduced the ‘scale-agnostic’ *G*_+_ discordance metric, however this internal metric is slow to calculate for large data. Furthermore, we show that *G*_+_ varies as a function of the proportion of observations in the predicted cluster labels (group balance), which is an undesirable property.

To address this problem, we propose a modification of *G*_+_, referred to as *H*_+_, and demonstrate that *H*_+_ does not vary as a function of group balance using a simulation study and with public single-cell RNA-sequencing data. Finally, we provide scalable approaches to estimate *H*_+_, which are available in the `fasthplus` R package.

## 1 Introduction

Quantifications of discordance such as Gamma (Goodman, Kruskal, 1979) and Tau (Kendall, 1938) have historically been derived to assess fitness from contingency tables. (The terms ‘discordance’ and ‘disconcordance’ have been used interchangeably to describe related metrics for contingency tables (Rohlf, 1974; Goodman, Kruskal, 1979), but in this work, we will use ‘discordance’). For all elements *α* ∈ *A* and *b* ∈ *B*, discordance is:

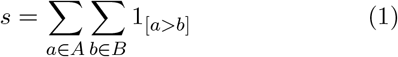

In this work, we consider a special case for *A* and *B* that explores the problem of unsupervised clustering (also known as observation partitioning). A typical clustering algorithm seeks to optimally group *n* observations into *k* groups (or clusters) using a dissimilarity matrix *D*(*n* × *n*) (e.g. Euclidean distance) or *d_ij_* for each *i, j* observations with 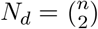 unique pairs.

If there does not exist a ground-truth label for each observation, internal validity metrics are used often to evaluate the performance of a set of predicted cluster labels *L* = [*l*_1_, …, *l_n_* : *l_i_* = 1 …, *k*] for a fixed *D*. Many internal fitness metrics quantify the tightness or consistency of partitions with functions such as within-cluster sums of squares (WCSS) or mean Silhouette scores (Rousseeuw, 1987). However, when comparing multiple dissimilarity measures, the interpretation of these performance metrics can be problematic as different dissimilarity measures have different magnitudes and ranges, leading to different ranges in the tightness of the clusters.

One solution is to use discordance as an internal validity metric that depends on the ranks of the dissimilarities, rather than on the dissimilarities themselves, thereby making it a ‘scale-agnostic’. For example, the discordance metric *G*_+_ (Williams, Clifford, 1971; Rohlf, 1974) uses the following to assess how well a given predicted cluster label *L* fits a dissimilarity *D* induced from the same observations (Rohlf, 1974; Desgraupes, 2018) (**Supplemental Note 1**):

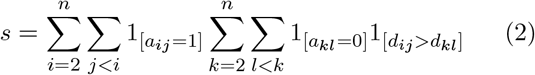

given fixed *L*, an adjacency matrix *A*(*n* × *n*) is defined using the predicted cluster label *l_i_*, *l_j_* for the *i*, *j* observations where *a_ij_* = 1 if *l_i_* = *l_j_* or *a_ij_* = 0 otherwise.

In the following sections, we first consider properties of *G*_+_ and demonstrate that *G*_+_ is slow to calculate for large data (due to the pairwise comparisons of dissimilarities in Equation 2). Furthermore, *G*_+_ (undesirably) varies as a function of the proportion of observations within the same predicted cluster labels (referred to as group or cluster balance). For example, when simulating ‘null’ data (random Gaussian data with no mean difference between two groups), the expected mean (and the interpretation) of the *G*_+_ discordance metric varies depending on the proportion of observations in each group (**Figure 1**). To ameliorate this, we propose a modification to *G*_+_, referred to as *H*_+_, and demonstrate that *H*_+_ does not vary as a function of group balance using a simulation study and with public single-cell RNA-sequencing (scRNA-seq) data. Finally, we provide scalable approaches to estimate *H*_+_, which are available in the `fasthplus` R package.

**Figure 1.**
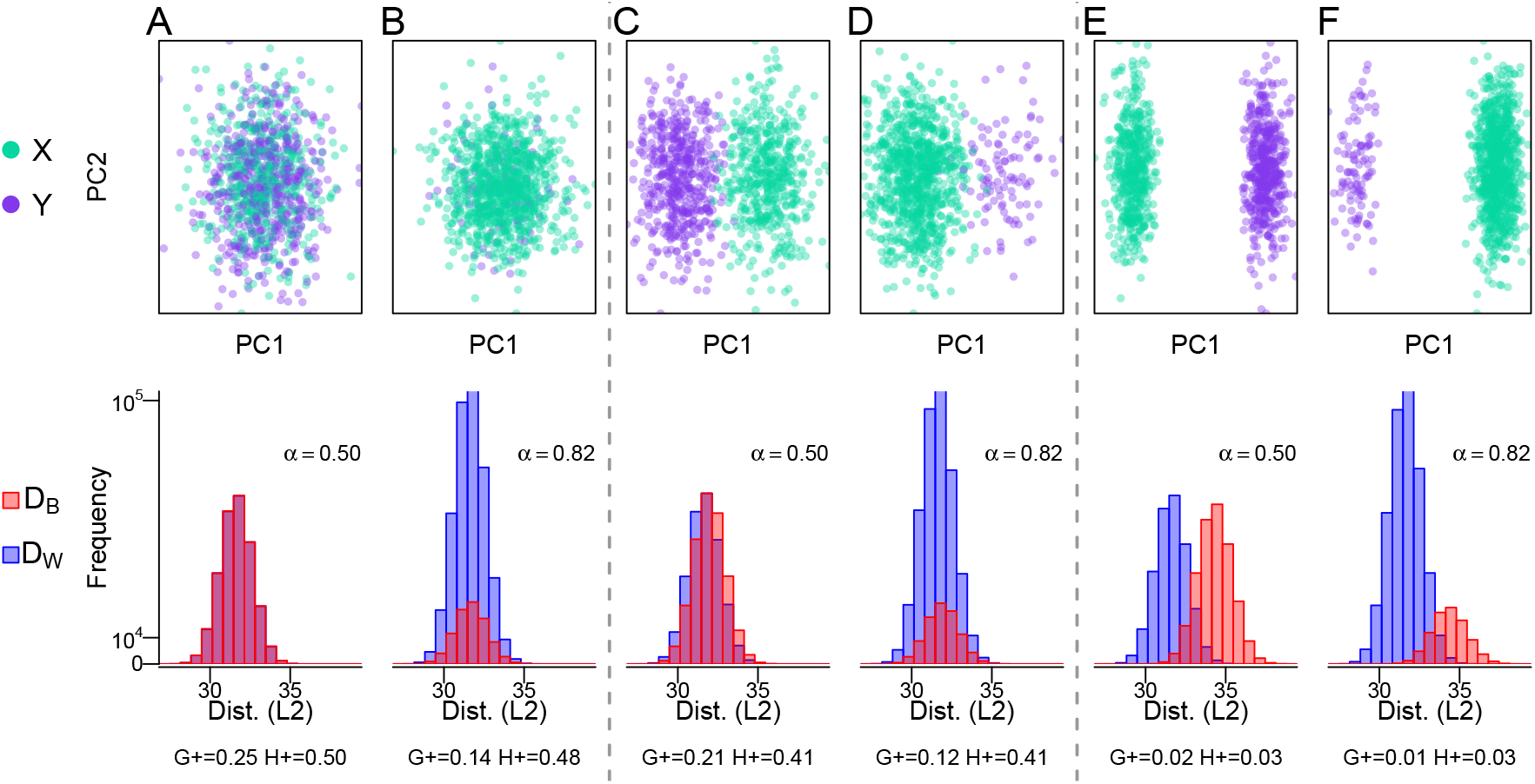
The *G*_+_ discordance metric varies as function of *α* (proportion of within-cluster distances), which is a function of the group balance |. We randomly sampled *n* = 1000 observations from a mixture distribution *b* * *X* + (1 – *b*) * *Y* with *b* being the probability of an observation coming from *X* ~ *N*(*μ_x_*, *σ*^2^) and 1 – *b* coming from *Y* ~ *N*(*μ_y_*, *σ*^2^) with **(A-B)** no mean difference (*μ_x_* – *μ_y_* = 0) (or a ‘null’ setting), **(C-D)** a small mean difference (*μ_x_* – *μ_y_* = 0.2), and **(E-F)** a large mean difference (*μ_x_* – *μ_y_* = 0.6). We simulate data with **(A,C,E)** balanced groups (*b* = 0.5) and **(B,D,F)** imbalanced groups (*b* = 0.9). For each simulation, the top row contains observation colored by group (*X* and *Y*) along the first two principal components (PCs) and the bottom row contains histograms of the within- (*D_W_*) and between- (*D_B_*) cluster distances (Euclidean) for the balanced and imbalanced groups. Refer to **Supplemental Figure S1** for an illustration of (and **Supplemental Note 2** for the explicit relationship between) the group balance and proportion of within-cluster distances, referred to as *α*. For each simulation, the bottom row includes *α* and the two discordance metrics *G*_+_ and *H*_+_. Generally, values close to zero represent more concordance, while a larger values represent more discordance.

## 2 The *G*_+_ discordance metric

The discordance metric *G*_+_ (Williams, Clifford, 1971; Rohlf, 1974) scales *s* (**Equation 2**) by 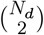, the number of ways to compare each unique distance to every other.

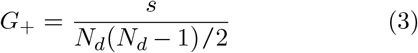

Generally, *G*_+_ close to zero represents two concordant sets, while a larger *G*_+_ is more discordant.

### 2.1 Applications of *G*_+_

The discordance metric *G*_+_ can be used to quantify the cluster fitness for given a dissimilarity matrix *D* and predicted cluster label *L*. For example, if *D* is fixed, lower values of *G*_+_ among many sets of labels *L*_1_, *L*_2_ … indicate increased cluster fitness (Rand, 1971; Williams, Clifford, 1971) or better performance. If we instead fix *L*, we can also use *G*_+_ to assess the fitness of multiple dissimilarity matrices *D*_1_, *D*_2_ … (Rohlf, 1974).

Because *G*_+_ depends on the relative rankings of pairwise distances, this transformation enables a robust comparison of dissimilarity measures by the structure they impose on the data rather than by the exact values of the distances themselves. This allows us to evaluate distances at highly varying scales without imposing bias with regard to the expected magnitude of distances. In other words, discordance metrics such as *G*_+_ offers a ‘scale-agnostic’ means to assess cluster fitness.

### 2.2 Properties of *s*

Consider *s* (**Equation 2**) with an adjacency matrix *A*(*n* × *n*) and dissimilarity matrix *D*(*n* × *n*). We denote the within-cluster distances as *D_W_* = {*d_ij_* : *a_ij_* = 1, *i* = 2, …, *n, j* < *i*} and the between-cluster distances as *D_B_* = {*d_kl_* : *a_kl_* = 0, *k* = 2, …, *n, l* < *k*}. The total number of distances in each of these sets (or length when vectorized) is |*D_W_* | and |*D_B_* |, respectively. As each upper triangular entry of *A* is binary (every distance is either between- or within-cluster), then |*D_W_*| + |*D_B_*| = *N_d_*.

We can define *α* where *α* ∈ (0, 1) as the proportion of total distances *N_d_* that are within-cluster distances. In this way, |*D_W_*| = *αN_d_*, and similarly, |*D_B_*| = (1 – *α*)*N_d_*. Then, conditional on *N_d_* and *α*, the *E*[*s*] is (**Supplemental Note 1**):

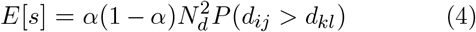

where *P*(*d_ij_* > *d_kl_*) is the probability that a within-cluster distance *d_ij_* ∈ *D_W_* is greater than a between-cluster distance *d_kl_* ∈ *D_B_*. This is the quantity we are interested in estimating, but there is a scaling factor 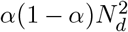 that depends on both *α* and *N_d_*. Next, we consider properties of first *P*(*d_ij_* > *d_kl_*) and then *G*_+_.

### 2.3 Properties of *P*(*d_ij_* > *d_kl_*)

If we know the expected mean and variance for *d_ij_* ∈ *D_W_* and *d_kl_* ∈ *D_B_*, we can estimate *P*(*d_ij_* > *d_kl_*). In the simple case where *E* [*D_W_*] = *E*[*D_B_*], we can consider *X* = *D_W_* – *D_B_*, then *E* [*X*] = 0 and a standardization of *X* demonstrates (assuming [co]variances exist) that 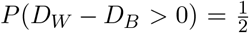. As we might expect, there is a 50% chance that *d_ij_* > *d_kl_* when *E*[*D_W_*] = *E*[*D_B_*].

### 2.4 Properties of *G*_+_

Using Equations 3 and 4, the expected value of *G*_+_ is (**Supplementary Note 1**):

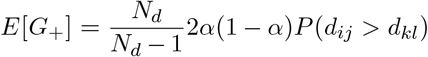

As 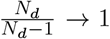 for large enough *N_d_*, then *E*[*G*_+_] has an undesirable property of being a function of 2*α*(1 – *α*), which has an explicit relationship with class balance *b* (**Supplemental Figure S1**, **Supplemental Note 2**).

For example, assume we randomly sampled *n* = 1000 observations from a mixture distribution *b* * *N*(*μ_x_*, *σ*^2^) + (1 – *b*) * *N*(*μ_y_, σ*^2^) with no mean difference (*μ_x_* – *μ_y_* = 0) and balanced classes (*b* = 0.5 and *α* = 0.5) then, we know 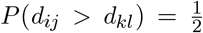 and 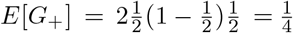. This can be thought of as a ‘null’ simulation where we expect no differences and *H*_+_ results in an intuitive value of 0.5, while *G*_+_ is 0.25. However, if there is an imbalance in class sizes (*b* = 0.9 and *α* = 0.82) then 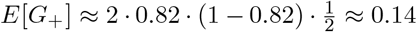. In practice, this is not a desirable property because the interpretation of *G*_+_ varies simply as a function of *α* (and class balance) (**Figure 1**).

## 3 The proposed method

### 3.1 An unbiased discordance metric with *H*_+_

To ameliorate this effect, we propose *H*_+_, which replaces the scaling factor *N_d_* (*N_d_* – 1)/2 in the denominator in *G*_+_ with 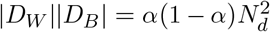:

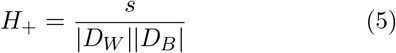

In other words, instead of scaling *s* by the total number of ways to compare every distance to every other distance, we divide by the number of ways to compare within-cluster distances to between-cluster distances. Hence, *E*[*H*_+_] is not a function of *α*:

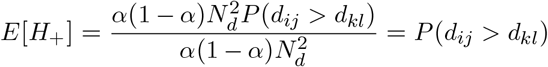

In fact, we can empirically verify that while *G*_+_ varies as a function of *α* (and hence class balance) (**Figure 2A**), *H*_+_ does not (**Figure 2B**), regardless of difference in expectation between the two groups.

**Figure 2.**
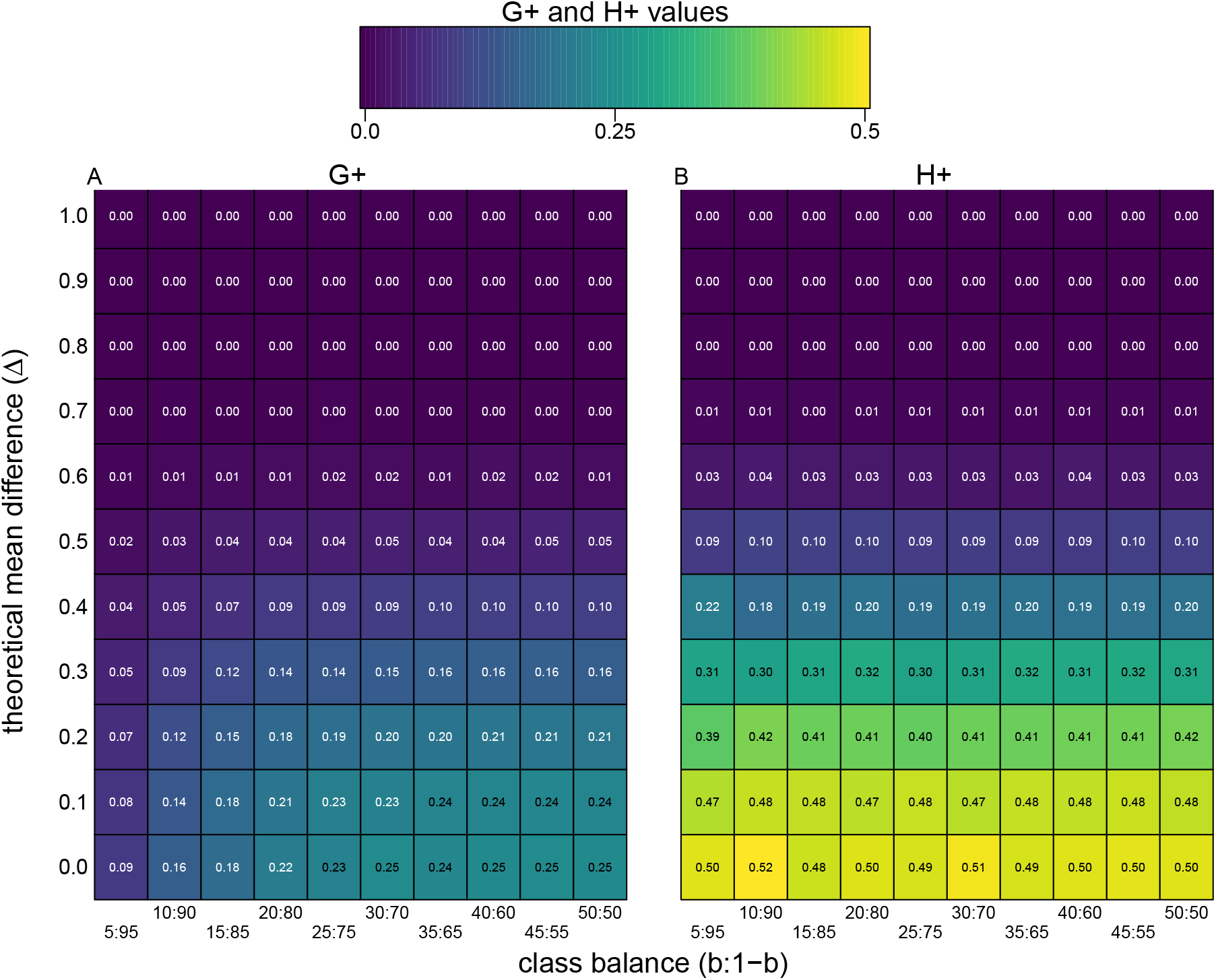
The *H*_+_ discordance metric does not change as a function of class balance |. We randomly sampled *n* = 1000 observations from a mixture distribution *b* * *X* + (1 – *b*) * *Y* with *b* being the probability of an observation coming from *X* ~ *N*(*μ_x_, σ*^2^) and 1 – *b* coming from *Y* ~ *N*(*μ_y_, σ*^2^) with a true mean difference (Δ = *μ_x_ – μ_y_*) (*y*-axis). Along the *x*-axis we change group (or class) balance from balanced (e.g. *b* = 0.50) and to imbalanced (e.g. *b* = 0.05) groups. The plots are heatmaps of true *G*_+_ (left) and *H*_+_ (right) discordance metrics, which shows *H*_+_ does not change as a function of class balance (*x*-axis), only as a function of the true effect size (*y*-axis).

### 3.2 Generalizing properties of *P*(*d_ij_* > *d_kl_*)

More generally, consider the function 1_[*d_ij_*>d_*kl*_]_. For some constant *c*, we can decompose this event as a joint event 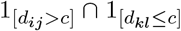 or 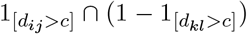 (Jardine, Sibson, 1968; Rohlf, 1974). Therefore, as *E*[*H*_+_] = *P*(*d_ij_* > *d_kl_*), we can decompose *E*[*H*_+_] into two quantities: *E*[*H*_+_] = *γ_W_γ_B_* where *γ_W_* = *P*(*d_ij_* > *c*) and *γ_B_* = (1 – *P*(*d_kl_* > *c*)). In other words, *H*_+_ empirically states a 100 × *γ_W_* % of *d_ij_* ∈ *D_W_* is strictly greater than 100 × *γ_B_* % of *d_kl_* ∈ *D_B_*. This implies *H*_+_ is not uniquely determined. For example, if *H*_+_ = 0.4, we could have *γ_W_* = 1.00, *γ_B_* = 0.4 or *γ_W_* = 0.80, *γ_B_* = 0.50. It should be noted that one can construct examples where two distinct pairs of *γ_W_*, *γ_B_* will have the same product, but do not imply each other.

### 3.3 Two algorithms to estimate *H*_+_ and *γ_W_*, *γ_B_*

One problem with the *H*_+_ (and *G*_+_) discordance metric (**Equation 5**) is that it requires the calculation of both (i) the dissimilarity matrix *D*(*n* × *n*) which scales *O*(*n^2^*) and (ii) *s* (**Equation 2**) which scales with the number of ways to compare within-cluster distances to between-cluster distances (or *O*(*n*^4^) comparisons). For example, with datasets of sizes *n* = 100 and 500, it takes 0.01 and 0.22 seconds, respectively, to calculate *D*(*n* × *n*) and it takes 0.08 and 59.68 seconds, respectively, to calculate *s* (**Figure 3A, Supplemental Table S1**). For datasets with more than *n*=500 observations, this quickly becomes computationally infeasible.

**Figure 3.**
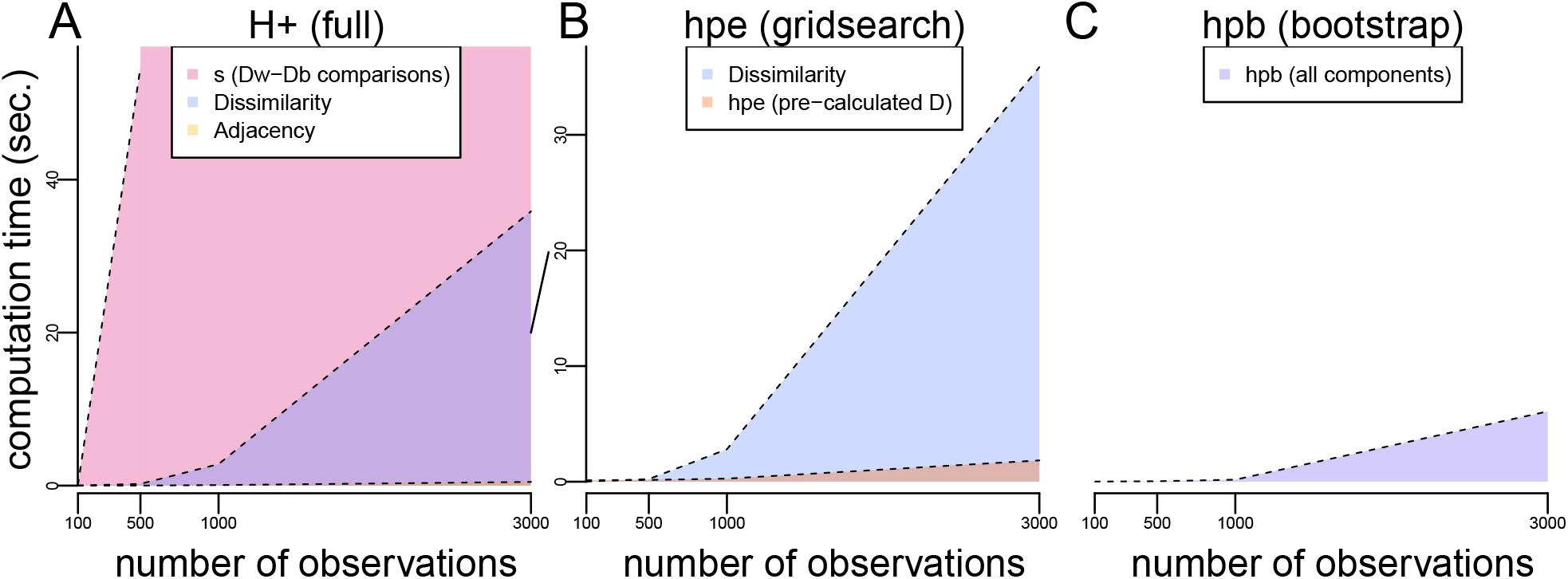
Computation times (seconds) for exact and approximate *H*_+_ calculations as a function of increasing number of observations *n* |. Computational time (*y*-axis) as a function of observations (*x*-axis) to calculate the individual components of **(A)** exact *H*_+_ including (i) the dissimilarity matrix *D*(*n* × *n*) (purple) scaling *O*(*n*^2^), (ii) the adjacency matrix (orange), and (iii) the most expensive operation *s* (pink) scaling *O*(*n*^4^). Note, *s* is only shown for *n*=100 and 500 observations, but the trend is shaded in for the other observations; **(B)** approximate *H*_+_ estimation (HPE) using the grid search procedure including (i) the dissimilarity matrix *D*(*n* × *n*) (blue) scaling *O*(*n*^2^) and (ii) the HPE algorithm to estimate *s* (orange) scaling *O*(*p);* **(C)** approximate *H*_+_ estimation using the bootstrap procedure (HPB) (purple), which scales similarly to HPE without the computational expense required for calculating *D*. Note (B) and (C) have a different *y*-axis scale than (A) for an zoomed in visualization of time.

To address this, we propose two algorithms to estimate *H*_+_ (an ‘h-plus estimator’ or HPE) inspired by the Top-Scoring Pair (TSP) (Leek, 2009; Magis, Price, 2012) algorithms, which use relative ranks to classify observations: a brute force approach with *O*(*p*^2^) comparisons, and a grid search approach with *O*(*p*) comparisons where *p* refers to percentiles of the data (rather than the *n* observations themselves) where *p* << *n*, leading to significant improvements in the computational speed to calculate *H*_+_. Specifically, both algorithms estimate *H*_+_ (referred to as *H_e_* or HPE) assume *D*(*n* × *n*) has been pre-calculated and provide faster ways to approximate *s* (**Figure 3B**). Both algorithms are implemented in `hpe()` function in the `fasthplus` R package.

Finally, in a later section (**Section 3.5**), we introduce a third algorithm based on bootstrap sampling to avoid calculating the full dissimilarity matrix *D*, thereby leading to further improvements in computational speed to estimate *H*_+_ (referred to as *H_b_* or HPB) (**Figure 3C**). The bootstrap algorithm is implemented in hpb() function in the `fasthplus` R package.

#### 3.3.1 Intuition behind HPE algorithms

The estimator *H_e_* (or HPE) assumes *D*(*n* × *n*) has been pre-calculated and then provides faster ways to approximate *s* (the pairwise comparisons of *D_W_* and *D_B_*). Specifically, we let the two sets *A* and *B* represent the ordered (ascending) dissimilarities *d_ij_* ∈ *D_W_* and *d_kl_* ∈ *D_B_*, respectively. Then, we bin the sets *A* and *B* into *p* + 1 percentiles where *q*(*A*)*_i_* and *q*(*B*)_*j*_ are the percentiles for *i, j* = 0, …, *p*. Note, *q*(*A*)_0_ = *min*(*A*) and *q*(*A*)_*p*_ = *max*(*A*). In both algorithms below, we check if *q*(*A*)_0_ > *q*(*B*)_*p*_, then *H_e_* = 1, and similarly, if *q*(*A*)_*p*_ < *q*(*B*)_0_ then *H_e_* = 0.

Next, we provide a graphical intuition for the two HPE algorithms by performing a simulation study. First, we simulate observations from two Gaussian distributions, namely *A* ~ *N*_10000_(0.3, 1) and *B* ~ *N*_10000_(−0.3, 1) and calculate the quantiles *q*(*A*) and *q*(*B*) for each of the sets with *p* + 1 = 11 (**Figure 4A**), *p* + 1 = 26 (**Figure 4B**), and *p* + 1 = 51 (**Figure 4C**). The calculation of these quantiles seeks to approximate the true ordered inequality information for each *d_ij_* ∈ *D_W_* and *d_kl_* ∈ *D_B_*. That is, if *D_W_*, *D_B_* were both given in ascending order, the white line in **Figure 4** shows the percent of *d_kl_* ∈ *D_B_* that is strictly less than each *d_ij_* ∈ *D_W_*. The true *H*_+_(≈ 0.66) is then given by the area under the white curve (the true rank orderings for each pair). Our goal is to use the following two algorithms to estimate the true *H*_+_ (fraction of blue area in the grid).

**Figure 4.**
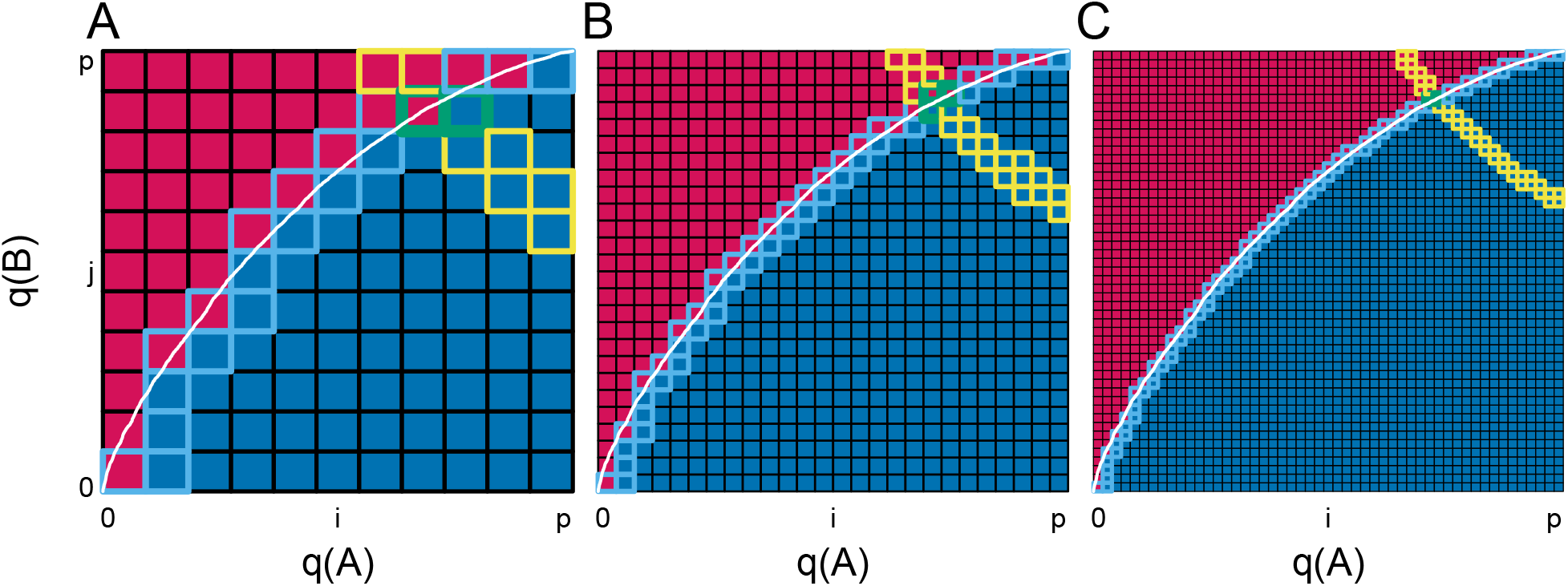
Graphical representation of two HPE algorithms to estimate *H*_+_ |. We simulate observations from two Gaussian distributions, namely *A* ~ *N*_10000_(0.3, 1) and *B* ~ *N*_10000_(–0.3, 1) and calculate the quantiles *q*(*A*) and *q*(*B*) for each of the sets with **(A)** *p* + 1 = 11, **(B)** *p* + 1 = 26, and **(C)** *p* + 1 = 51. The white curve represents the percent of elements in *A* that are strictly less than each element in *B*. The goal is to estimate the true *H*_+_(≈ 0.66) (area under the white curve) using one of two HPE algorithms. The brute force approach (**HPE algorithm 1**) uses Riemann integration to approximate the white curve by summing the area of the blue squares below the curve. The grid search approach (**HPE algorithm 2**) starts at the minimum of *q*(*A*) and *q*(*B*) and moves along the red-blue border to approximate the white curve (path followed represents the squares with the light blue borders). The HPE contour *H_e_* (or estimate of *H*_+_) 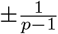 is given by yellow-bordered squares. In other words, every pair *γ_W_*, *γ_B_* such that 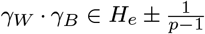, the interval guaranteed to contain *H*_+_). The intersection of this yellow contour 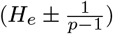 and blue contour (grids visited by HPE algorithm 2) are the green-bordered squares, which represents the numerical estimate for *γ_W_* and *γ_B_*.

#### 3.3.2 HPE algorithm 1: *H*_+_ (brute force)

Algorithm 1 numerically approximates *H*_+_ with Riemann integration. Specifically, using a double `for` loop with (*p* + 1)^2^ comparisons, this brute force approach sums the area of the squares that are blue, resulting in an algorithm on the order of *O*(*p*^2^). The path taken by our implementation of this algorithm is given by the squares with light blue borders, and the contour corresponding to the true *H*_+_ ≈ .66 is (approximately) represented by the squares with yellow outlines (**Figure 4**).

##### Algorithm 1 *H*_+_ (brute force)

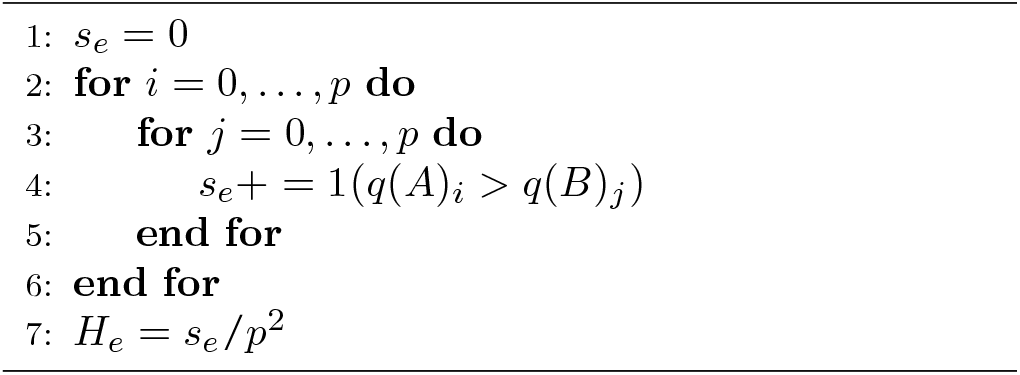

#### 3.3.3 HPE algorithm 2: *H*_+_ (grid search)

An alternative and faster approach (on the order of *O*(*p*) comparisons) is to sketch the surface (blue-red border) that defines *H*_+_. By starting at the minimum of *q*(*A*) and *q*(*B*), Algorithm 2 moves along the blue-red border that defines *H*_+_ using grid search to determine whether to increase *q*(*A*)*_i_* or *q*(*B*)*_j_* with each iteration.

##### Algorithm 2 *H*_+_ (grid search)

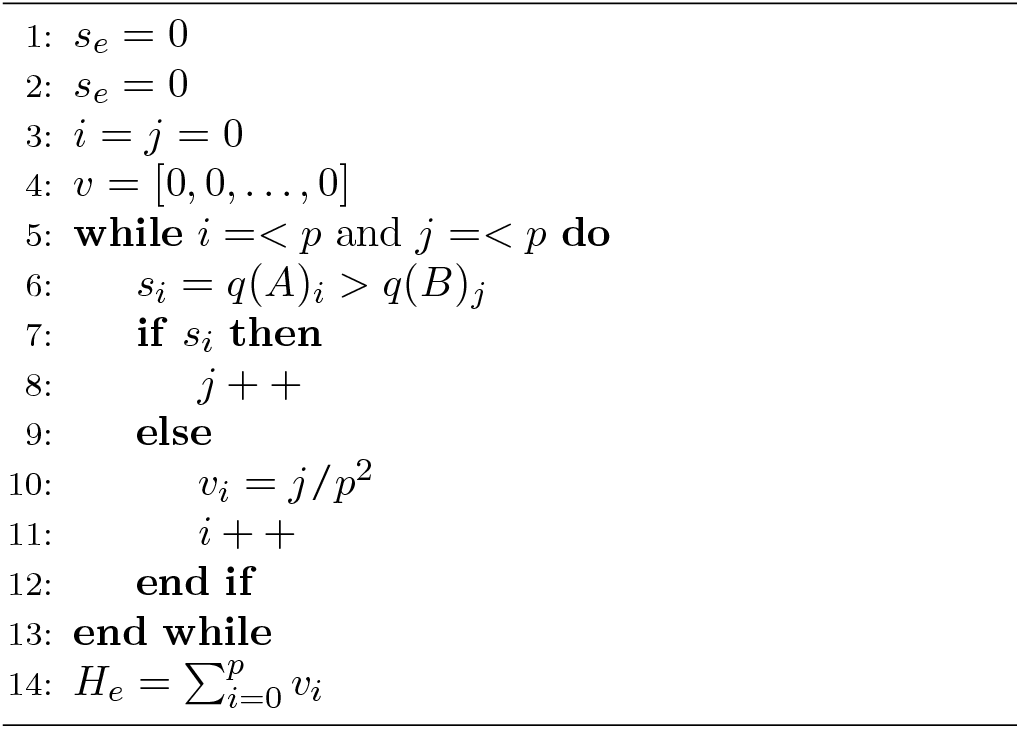

### 3.4 Convergence of HPE algorithms 1 and 2

Next, we provide a numerical bound for the accuracy of *H_e_* for both the brute force and grid search approaches. For each *q*(*A*)_*i*_, *i* = 0, …, *p*, HPE algorithm 2 (and intrinsically algorithm 1) ascertains one of the following:

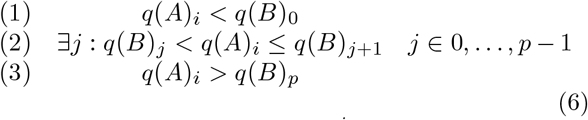

In (1), we have confirmed that 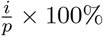 of *A* are less than 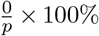 of *B* and the *i^th^* addition to the numerical integral will be zero, that is, *v_i_* = 0 in HPE algorithm 2. In (3), we see that 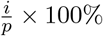 of *A* are greater than 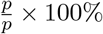 of *B* and 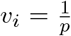 in HPE algorithm 2. In (2), we know that 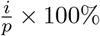 of *A* are bigger than 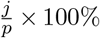 of *B*, but not greater than 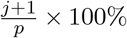 of *B*, and 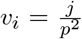 in HPE algorithm 2. Recall that *H_e_* is estimated as the sum over each *v_i_* where 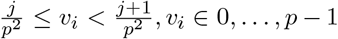. We denote *u_i_* as the true value of this sum for column *i*, that is, 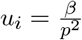 where 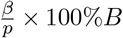 are less than or equal to *q*(*A*)*_i_* and 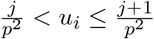. Thus, for (2), we have the condition 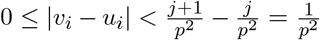, in other words, the addition to *H_e_* from the *i^th^* column will differs from the true value (*H*_+_) by at most 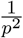. Thus, for all *p*:

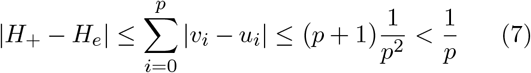

That is, by taking *p* + 1 percentiles of *D_W_* and *D_B_*, our estimate for HPE algorithm 2 will be within 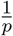 of *H*_+_. This follows when one considers HPE algorithms 1 and 2 are approximations of the paired true rank comparisons (white curve in **Figure 4**) using Riemann integration with increasing accuracy as a function of *p*. An additional argument for the convergence of these algorithms is presented in **Supplemental Note 3**.

#### 3.4.1 Estimating *γ_W_* and *γ_B_*

To estimate *γ_W_* and *γ_B_*, we use the intersection of the yellow contour 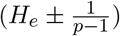 and blue contour (path visited by HPE algorithm 2), which are the green-bordered squares in **Figure 4**. As our approach guarantees that 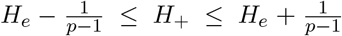, we can identify every pair 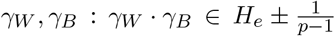 as potential values of *γ_W_*, *γ_B_* : *γ_W_* · *γ_B_* ≈ *H*_+_. Our algorithm also identifies the values of *γ_W_*, *γ_B_* that are true for the observed data (all area below the white line in **Figure 4**) or those which have been verified as false (all area above the white line in **Figure 4**). Our estimate for *γ_W_*, *γ_B_* is then the intersection of *γ_W_*, *γ_B_* that are empirically verified in HPE algorithm 2 such that 100 × *γ_W_* % of *d_ij_* ∈ *D_W_* (*γ_W_* ∈ [0, 1]) is strictly greater than 100 × *γ_B_* % of *d_kl_* ∈ *D_B_* (blue squares in **Figure 4**) and *γ_W_*, *γ_B_* that satisfy *γ_W_ · γ_B_ ≈ H*_+_ (yellow squares in **Figure 4**).

### 3.5 Bootstrap algorithm to estimate *H*_+_

As noted in Section 3.3, while the computational speed of the HPE algorithms for identifying ways to approximate *s* are significantly faster than calculating the full *H*_+_ (**Figure 3A-B**), both of these algorithms assume the dissimilarity matrix *D*(*n* × *n*) has been pre-computed and that an adjacency matrix *A*(*n* × *n*) must be calculated. Unfortunately, the *O*(*n*^2^) computational requirements for full pairwise dissimilarity calculation to quickly becomes infeasible.

To address the limitation of computing and storing all pairwise dissimilarities, we implemented a bootstrap approximation of *H*_+_ (HPB or *H_b_*) that samples with replacement from the original *n* observations *r* times (bootstraps) with a per-bootstrap sample size *t*. We sample proportionally according to the vector *b* as described in **Supplemental Note 2**, that is, each of the *k* clusters is randomly sampled *t_j_* ≈ *b_j_* × *t* times where 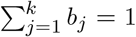 such that 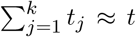. For each of *r* iterations, the *t* sampled observations are used to generate dissimilarity and adjacency matrices which are then used to calculate a point estimate of *H*_+_. The mean over these *r* bootstraps is *H_b_*, the bootstrap *H*_+_ estimate. The bootstrap approach scales substantially better than full dissimilarity calculation (**Figure 3C**). In our simulations, bootstrap parameters *r* = 0.05 × *n*, *t* = 100 yield *H*_+_ estimates within 0.01 of that given by HPB with *p* = 10001 (1*e* – 04 accuracy) with economical performance improvements (for example, a reduction in computation time from 38.57 with HPE to 5.74 with HPB at 3000 observations) (**Supplemental Figure S3 and Table S1**).

## 4 Application of *H*_+_ to the analysis single-cell RNA-sequencing data

In this section, we demonstrate the use of *H*_+_ as an internal validity metric in application of scRNA-seq data with predicted cluster labels. Also, we compare *H*_+_ to other widely-used validity measures, including both (i) external (i.e., comparing predicted labels to ground-truth clustering known *a priori*) and (ii) internal (derived from the data itself) measures (Halkidi et al., 2001; Theodoridis, Koutroumbas, 2009).

### 4.1 Motivation

Consider a scRNA-seq dataset with *n* observations (or cells) each with *G* features (or genes). We introduced and formulated *H*_+_ an internal validity metric to assess the fitness of a single dissimilarity measure *D* and label *L*. Here, we introduce two scenarios where the goal is to compare the performance of either (i) two sets labels *L_m_*, *L*_*m*–1_ and a fixed dissimilarity *D*, or (ii) two dissimilarity measures *D_m_*, *D*_*m*–1_ with a fixed label *L*. In first scenario, *m* and *m* – 1 could represent two iterations in a single clustering algorithm or they could be labels from two separate clustering algorithms. As *E*[*H*_+, *m*_] = *P*(*d_ij_* > *d_kl_*)_*m*_ (and similarly with *m* – 1), the condition *H*_+, *m*_ < *H*_+, *m*–1_ can be re-written as follows

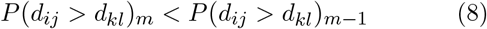

As *N_d_* is fixed in the following subsections, we offer interpretations of the condition in Equation 8 for fixed *L* with varying *D* and fixed *D* with varying *L*.

### 4.2 Data

We used the `sc_mixology` (Tian et al., 2019) scRNA-seq dataset, which provides an experimentally derived “gold standard” true cell type identity (label) for each cell (https://github.com/LuyiTian/sc_mixology/).

The UMI counts and cellular identities were obtained for *n* = 902 cells comprised of three cell lines (H1975, H2228, and HCC827). The cell lines are used as the true cell type labels. Raw counts were log 2-normalized with a pseudocount of 1, and per-gene variance was calculated using `scran` (Lun et al., 2016). Dissimilarities (L1 and L2) were calculated using log 2-normalized counts and the top 1000 most variant genes. Dendrograms were induced directly from the L2 dissimilarity using complete and weighted-average linkages. Cluster labels were induced by cutting each dendrogram at the true value of *k* = 3.

### 4.3 Fixed *L* varying *D*

If a user were generating an analysis pipeline, prior to deployment, it may be insightful to compare the performance of several dissimilarity measures on a previously validated label-dataset pair (Baker et al., 2021). In this case, fixing *L* will imply that *α_m_* = *α*_*m*–1_, then from Equation 8 we know that *s_m_* < *s*_*m*–1_ for two dissimilarity matrices *D_m_* and *D*_*m*–1_. That is, the number of within-cluster distances greater than between-cluster distances will have strictly decreased.

### 4.4 Fixed *D* varying *L*

Similarly, *D* can be fixed (e.g. Euclidean distance) with a goal to compare the fitness of one generated label set *L_m_* (i.e., iteration *m* of a clustering algorithm) to a previous label *L*_*m*–1_. Also we note from Equation 8, this implies while *α_m_* ≠ *α*_*m*–1_, then the discordance has decreased. To demonstrate the use of *H*_+_ as a cluster fitness metric, we first induce labels from four hierarchical clustering methods (**Figure 5A-D**), and compare against well-known both external and internal validity metric (**Figure 5E-F**).

**Figure 5.**
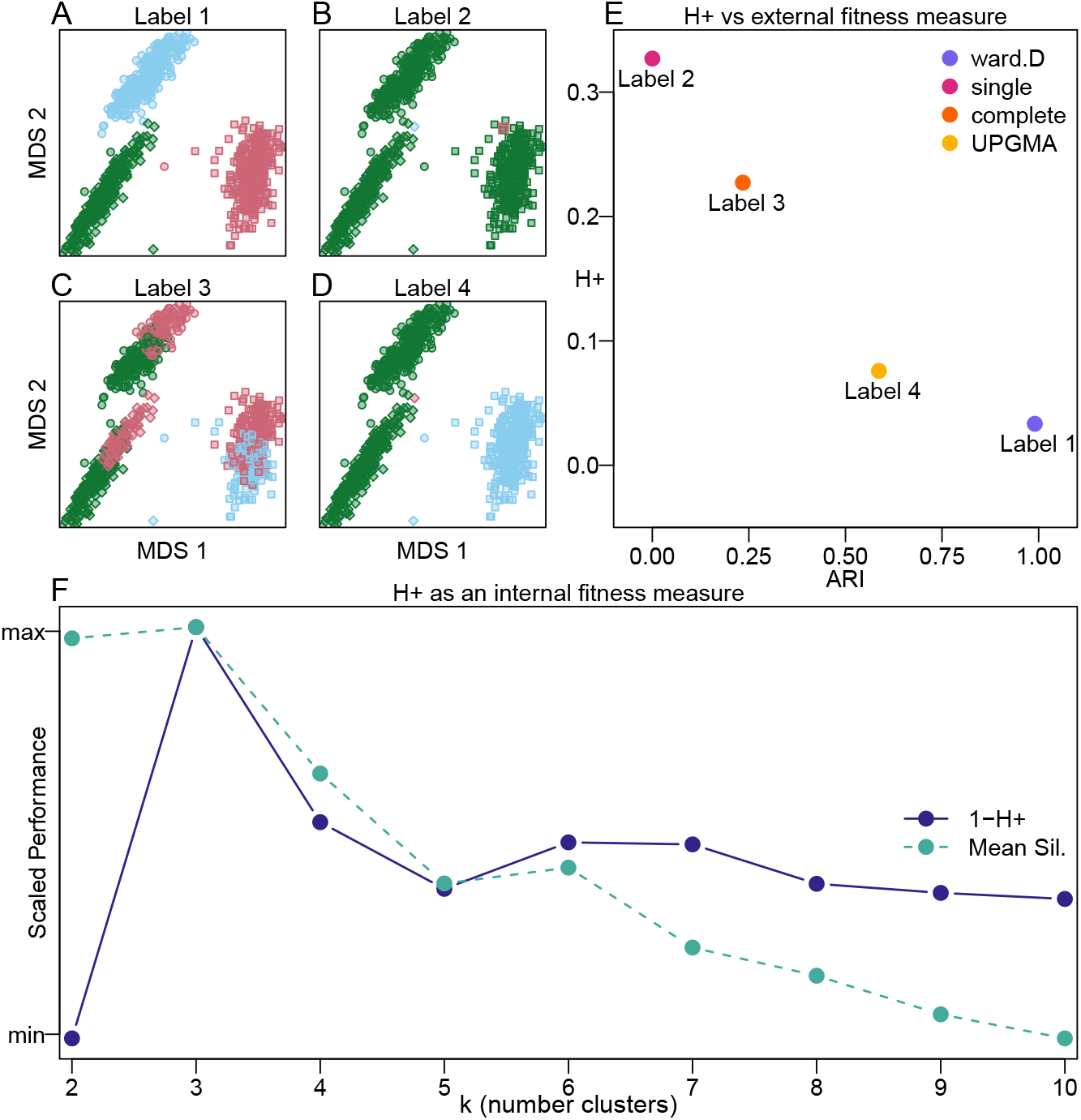
The *H*_+_ metric is an internal validity measure for assessing the performance of induced cluster labels |. Multidimensional scaling (MDS) plots with shapes representing true cell type labels from the `sc_mixology` scRNA-seq dataset and colors representing induced (or predicted) cluster labels from four hierarchical clustering methods implemented in the `hclust()` function in the base R `stats` package including **(A)** Ward’s method, **(B)** single linkage method **(C)** complete linkage method, and **(D)** unweighted pair group method with arithmetic mean (UPGMA). **(E)** Scatter plot of *H*_+_ (an internal validity metric) compared to Adjusted Rand Index (ARI) (an external validity metric) demonstrating shared information between the two metrics, which *H*_+_. recovers without the need of an externally labeled set of observations. **(F)** An “elbow” plot comparison between two internal validity metrics (*y*-axis scaled between 0 and 1): (i) mean silhouette score and (ii) 1 – *H*_+_. (for ease of comparison) calculated from labels induced using with *k* = 2, …, 10 (*x*-axis). The peak of both metrics at the correct *k* = 3 indicates that *H*_+_ accurately identifies the most accurate label set when compared to a second established internal fitness measure.

First, we compare *H*_+_ as an internal validity metric to an external validity metric, namely the Adjusted Rand Index (ARI), which assess the performance of the induced cluster labels using a gold-standard set of cell type labels in the `sc_mixology` (Tian et al., 2019) scRNA-seq dataset. Here, the induced labels with better (higher) ARI also posses better (less) *H*_+_ discordance (**Figure 5E**). In this sense, *H*_+_ (an internal validity measure without the dependency of a gold-standard set of labels) captures similar information as ARI (an external validity measure that depends on the use of a gold-standard set of labels).

Next, we compare *H*_+_ as an internal validity measure to other internal validity measures. Specifically, we induce labels using partition around medoids (*k*-medoids clustering) for values of *k* = 2, …, 10. For each label and *k*, the mean Silhouette score (Rousseeuw, 1987) and *H*_+_ were calculated. We found that *H*_+_ accurately identifies the correct *k* = 3 for induced labels when compared to an internal validity metric (that is, how well the data are explained by a single set of labels) (**Figure 5F**). The within-cluster sum of square dissimi-larities (**Supplemental Figure S2**), and indicates the same trend as *H*_+_ and the Silhouette score.

## 5 Discussion

Quantifying how well a generated clustering fits the observed data is an essential problem in the statistical and computational sciences. Most methods for measuring cluster fitness are explicitly valued on the dissimilarity induced from the data. While appealing in their simplicity and interpretation, these approaches are potentially more susceptible to numerical bias between observations or type of dissimilarity measure. Discordance metrics, such as *τ* and *G*_+_ circumvent this issue by assessing label-dissimilarity fitness implicitly on the dissimilarity values. In this work, we show *G*_+_ is as an estimator for the probability that a within-cluster dissimilarity is strictly greater than a between-cluster dissimilarity, *P*(*d_ij_*) > *P*(*d_kl_*). However, we also show that *G*_+_ varies as a function of the proportion of within-cluster distances (*α*), which an undesirable property of the discordance metric.

Here, we present *H*_+_, a modification of *G*_+_ that retains the scale-agnostic discordance quantification while addressing problems with *G*_+_. Explicitly, *H*_+_ is an unbiased estimator for *P*(*d_ij_*) > *P*(*d_kl_*). This benefit is most easily seen in the manner that *H*_+_ will be unaffected by the value of *α*, the balance of between- to within-cluster distances, a formulation which permits the user to assess fitness for arbitrary value of *k*. We discuss theoretical properties of this estimator, provide two simple algorithms for implementation, and ascertain a strict numerical bound for their accuracy as a function of a simple user-defined parameter. We also introduce an estimate of *H*_+_ based on bootstrap resampling from the original observations that does not require the full dissimilarity and adjacency matrices to be calculated.

As *H*_+_ can be used to assess the fitness of multiple dissimilarities for a fixed label, or to compare multiple labels given a fixed dissimilarity, we envision that *H*_+_ can be employed in both development and analysis settings. If the true observation identities (labels) are known for a dataset, *H*_+_ could be utilized in the development stages of analytical software and pipelines to ascertain the most advantageous dissimilarity measure for that specific problem. In the alternate setting, we envision that *H*_+_ can be used to quantify performance in clustering/classification scenarios. If the true labels are unknown, *H*_+_ could be used to identify the most robust clustering algorithm for a fixed dissimilarity measure. As a possible future direction, one could imagine directly minimizing discordance as the objective criteria within a clustering algorithm for optimizing iterative labels. Intuitively, a sequence of labels with decreasing discordance indicates that the generated labels have more accurately described the dissimilarity structure of the data.

Due to its generalizability to the number of clusters *k* or the balance of within to between-cluster dissimilarity *α*, *H*_+_ may be susceptible to degenerate cluster labels. For example, in the hierarchical clustering portion of **Figure 5**, Label 4 is less discordant than Label 3 in terms of both *H*_+_ and ARI. Label 4 has simply merged two true clusters, and placed a single point in a third identity. While Label 4 is more accurate than Label 3, it achieves this by exploiting an opportunity to increase the proportion of same-cluster pairs, that is, maximizing *α*. One could also imagine a scenario where an algorithm simply makes *k* very large to minimize *α*. In both scenarios, the labels generated are unlikely to be particularly informative for the user. We posit that some form of penalization for *H*_+_ may help to alleviate these degenerate cases. For example, dividing *H*_+_ by `max`{*α*, 1 – *α*} is a penalty for degeneracy in the case of putting many obervations in the same label. Conversely, a division by `min`{*α*, 1 – *α*} is a potential penalty for the other degeneracy of making many very small clusters.

We also imagine that discordance measures can be synthesized with probabilistic dissimilarity frameworks such as locality-sensitive hashing (LSH) and coresets (Datar et al., 2004; Har-Peled, Mazumdar, 2004). For example, it could be useful if theoretical (probabilistic) guarantees of observation proximity from LSH algorithms could be extended to similar guarantees for the discordance of observations embedded in the hash space. It may also prove fruitful to explore discordance outside the scope of the clustering/classification problem, such as pseudotime (1-dimensional ordering) or ‘soft’ (weighting membership estimation) clustering problems.

In practice, *H*_+_ could provide an additional means to consider termination of a clustering algorithm in a distance-agnostic manner. For example, the *k*-means algorithm (Hartigan, Wong, 1979) and its variants seek to minimize a form of the total within-cluster dispersion (dissimilarity). These and algorithms with similar objective functions are subject to changes in behavior as the distance function changes. The extent to which minimizing discordance such as *H*_+_ provides benefits regarding sensitivity to noise and magnitude of the distances is intriguing and outside the scope of this work.

## Code and software availability

All analyses and simulations were conducted in the `R` programming language. Code for reproduction of all plots in this manuscript is available at https://github.com/stephaniehicks/fasthpluspaper.

Both HPE and HPB have been implemented in the `fasthplus` package in `R` available via GitHub https://github.com/ntdyjack/fasthplus (submitted to `CRAN`).

## Author Contributions

ND and SCH developed the *H*_+_ method and the `fasthplus` R package. ND performed both simulation studies and single-cell gene expression analyses. ND and SCH wrote the manuscript with contributions from DB, VB, BL. All authors read and approved the final manuscript.

## Acknowledgements

The authors would like to thank Kasper Hansen for the pre-print template and the Joint High Performance Computing Exchange (JHPCE) for providing computing resources.

## Supplementary Materials

### Supplemental Notes

#### Note 1

Assume we have a set of *n* unique observations. For a given dissimilarity matrix *D* (e.g. Euclidean distance):

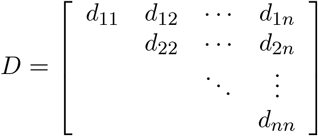

and fixed set of predicted cluster labels *L*, we can generate an adjacency matrix that tells us whether each observation has the same label (i.e., falls in the same cluster)

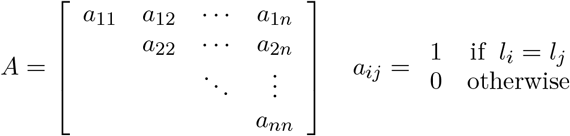

Using *d_ij_* and *a_ij_* for each *i*, *j* pairs of observations, we can then rewrite *s* in Equation (1) in terms of *A*, *D*. Let *A* = *D_W_* = {*d_ij_* : *a_ij_* = 1; *i* = 2, …, *n*, *j* < *i*}, that is, looping over the upper-triangular elements of *A*, *D* such that *a_ij_* = 1 to select pairs which correspond to observations that are within the same cluster. We similarly define *B* = *D_B_* = {*d_kl_* : *a_kl_* = 0, *k* = 2, …, *n*, *l* < *k*}, the dissimilarity pairs corresponding to observations with different cluster labels.

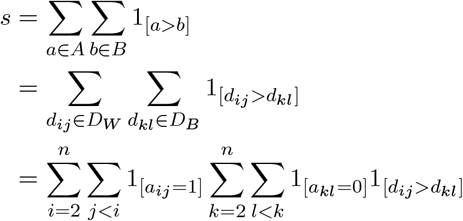

The total number of distances in each of these sets is |*D_W_*| and |*D_B_*|, respectively. As each upper triangular entry of *A* is binary (every distance is either between or within-cluster), we know that |*D_W_*| + |*D_B_*| = *N_d_*. We can define *α* where *α* ∈ (0, 1) as the portion of total distances *N_d_* that are within-cluster distances (or *d_ij_* ∈ *D_W_*). In this way, we can define |*D_W_*| = *αN_d_*, and similarly, |*D_B_*| = (1 – *α*)*N_d_*.

Conditional on *N_d_* and *α*, the expected value of *s* is

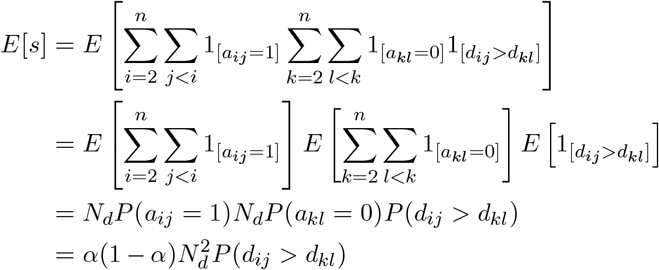

where *P*(*d_ij_* > *d_kl_*) is the probability a within-cluster distance *d_ij_* ∈ *D_w_* is greater than a between-cluster distance *d_kl_* ∈ *D_B_*.

Then, the expectation of *G*_+_ is:

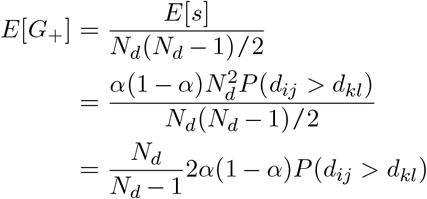

#### Note 2

The relationship between *α* (the portion of unique dissimilarities that correspond to pairs of observations in the same cluster) and the balance (portion of cells assigned to each of *k* groups) of these clusters is discussed below. Using the notation above for and an arbitrary label *L* (a vector of length *n*, *L_i_* ∈ 1, …, *k*) where *L_i_* = *j* indicates that the *i^th^* observation is assigned membership to the *j^th^* cluster group. We define the portion of observations in group *j* as follows

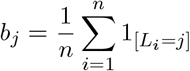

By definition we will have that 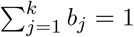. Each of the *k* clusters will contribute to the quantity *α*, a fraction of the *n*(*n* – 1)/2 unique pairs. It can be seen that for the *j^th^* cluster, this contribution is the upper triangular elements of a matrix block with size *n* · *b_j_*

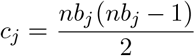

We now express *α* as a sum over each of *k* contributions *c_j_*, *j* = 1, …, *k*

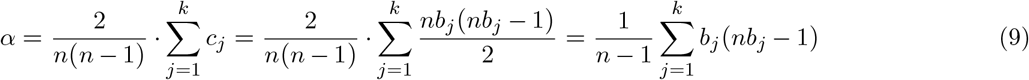

#### Note 3

We can also consider convergence in terms of the sum s by considering *q*(*D_W_*) and *q*(*D_B_*) as sampling without replacement from *D_W_* and *D_B_*. We denote *s_e_* the estimated form of the sum *s* Eq(2), that is

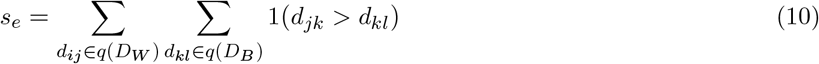

While |*q*(*D_W_*)| ≤ |*D_W_*| and |*q*(*D_B_*)| ≤ |*D_B_*|, we have that *s_e_* + *s_n_* = *s* where *s_n_* represents portions of the summand *s* that have not yet been counted in *s_e_*. This allows us to consider the convergence of an estimated *H_e_* to the true *H_e_* in terms of the decomposition *s_e_* = *s* – *s_n_*

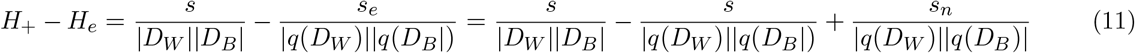

The denominators of tthe second and third term approach |*D_W_*||*D_B_*| as the number of distances sampled increases. The second term seems to approach H+ at 1/|*q*(*D_W_*)||*q*(*D_B_*)|. The third term approaches zero as |*q*(*D_W_*)||*q*(*D_B_*)| → |*D_W_*||*D_B_*| and *s_n_* decreases with each iteration in a factor bounded by |*q*(*D_W_*)||*q*(*D_B_*)|. This argument provides an intuitive argument that the convergence if is achieved simply by increasing |*q*(*D_W_*)| and |*q*(*D_B_*)|.

### Supplemental Figures

**Supplementary Figure S1.**
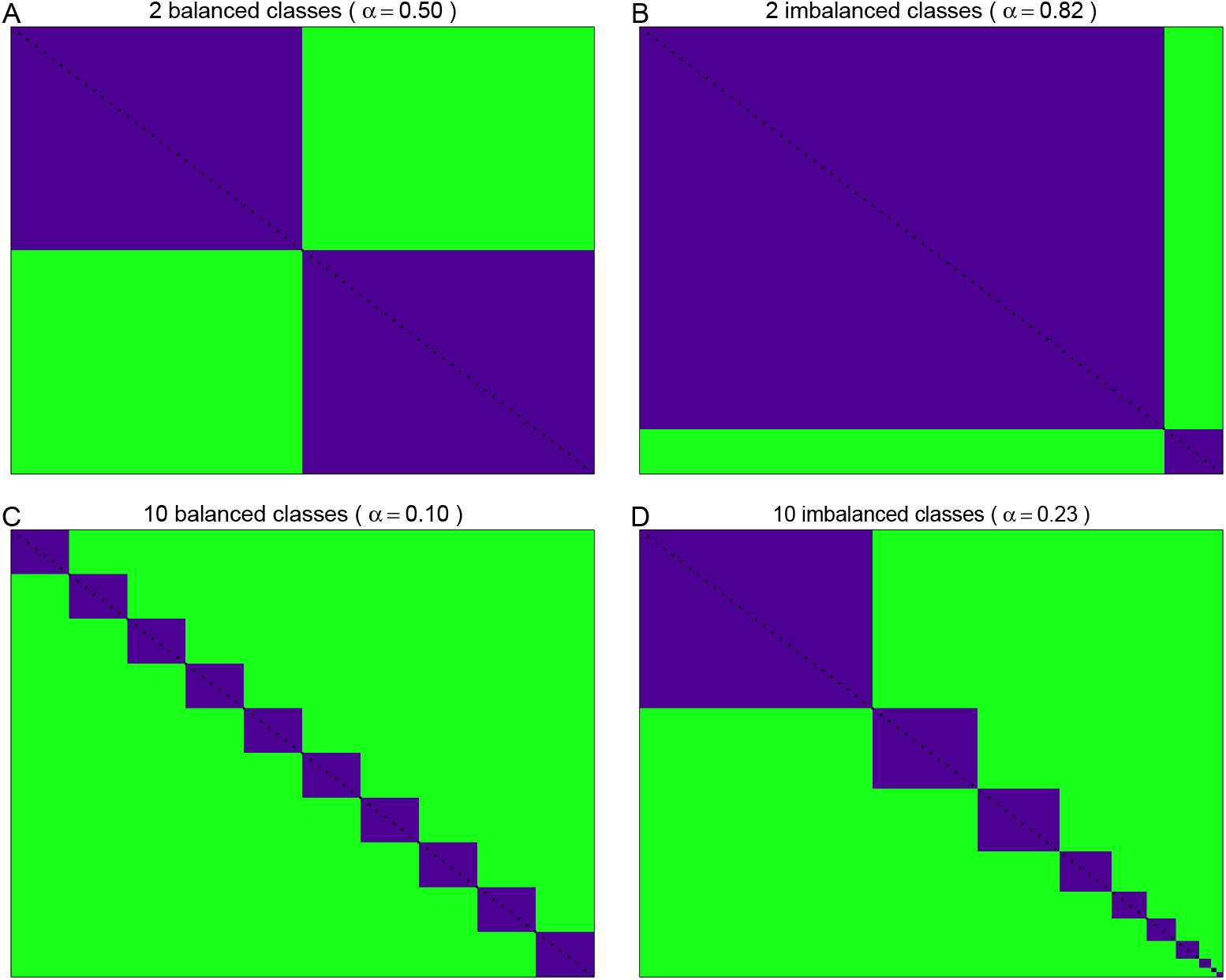
An illustration of the relationship between group (or class) balance and *α* |. Recall that *α* is the portion of total distances *N_d_* that are within-cluster distances, or |*D_W_* | = *αN_d_*, which can be seen most easily by using a heatmap of an adjacency matrix for **(A-B)** two or **(C-D)** ten groups for **(A,C)** balanced or **(B,D)** imbalanced groups. Unique pairs of observations are given on the upper-triangular potion of each adjacency matrix. Within-cluster pairs are given in blue, and between-cluster pairs are given in green. In other words, *α* is given by the blue proportion of each adjacency matrix. The group balance (and corresponding *α*) is **(A)** *b*_1_, *b*_2_ = 0.5 (*α* = 0.50), **(B)** *b*_1_ = 0.9, *b*_2_ = 0.1 (*α* = 0.82), **(C)** *b*_1_, …, *b*_10_ = 0.1 (*α* = 0.10), and **(D)** *b*_1_, *b*_2_, …, *b*_10_ = 0.40, 0.18, 0.14, 0.09, 0.06, 0.05, 0.04, 0.02, 0.01, 0.01 (*α* = 0.23).

**Supplementary Figure S2.**
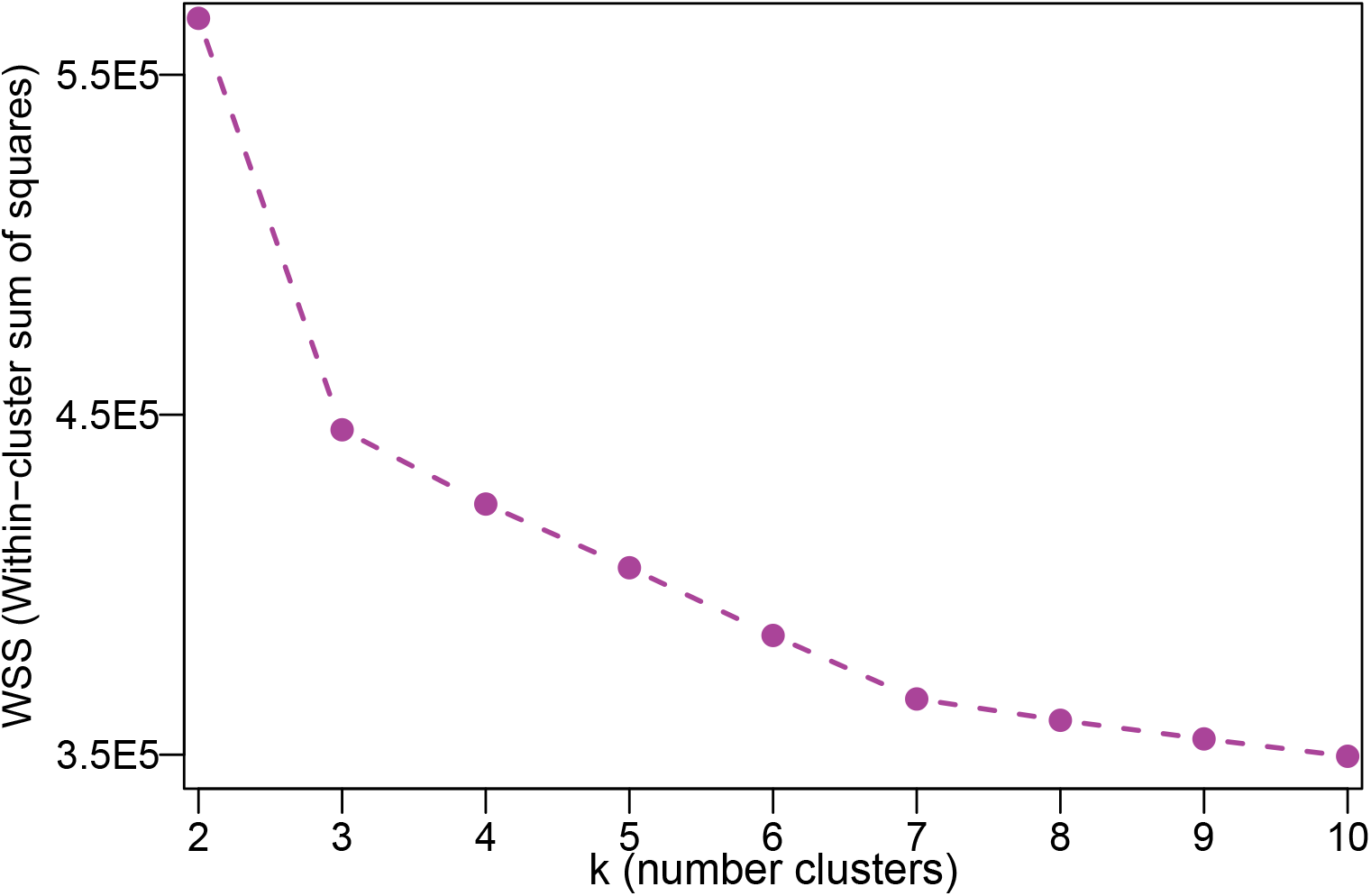
Within sum of squares (WSS) “elbow” plot values for k-medoids clustering in Figure 5 |. The sum of square within-cluster dissimilarities as a function of *k* = 2, …, 10 is a practical heuristic for estimating the correct *k* in many clustering algorithms. The analyst seeks to identify the “elbow” of this function, that is, smallest value of *k* at which in *WSS*(*k*) ceases to rapidly decrease for greater *k*. As in Fig5, the elbow at *k* = 3 indicates that the most accurate label set is likely given by this *k*.

**Supplementary Figure S3.**
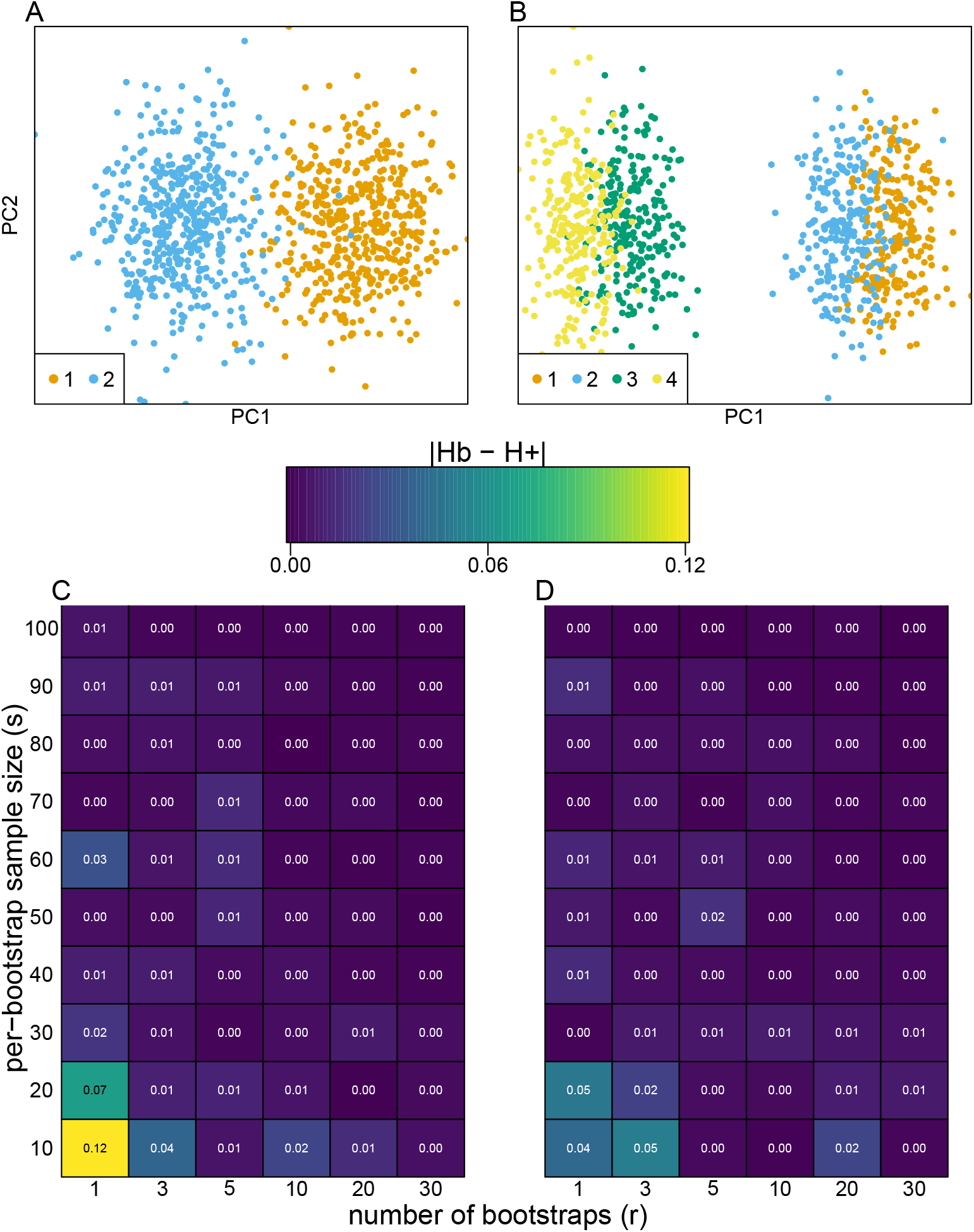
Accuracy of the bootstrap *H*_+_ estimation procedure for two simulated datasets |. Two data were simulated using 1000 observations, 500 features with two (*N*(–0.05, 0.25), *N*(0.05, 0.25)) **(A)** and four (*N*(–0.15, 0.25), *N*(–0.10, 0.25), *N*(0.10, 0.25), *N*(0.15, 0.25)) **(B)** balanced classes. The difference absolute difference between *H*_+_ (estimated using HPE with *p* = 10001) and the bootstrap estimate *H_b_*, for *r* replications (bootstraps) using *s* samples per bootstrap. For these simulations, sampling as little as 1% (*t* = 10) of the observations over *r* = 30 bootstraps provides an accurate estimate for *H*_+_.

### Supplemental Tables

**Supplementary Table S1.** Performance evaluation for elapsed time as reported in Figure 3 |. We report the elapsed time (seconds) for the individual components including calculating the dissimilarity matrix (dis), the adjacency matrix (adj), *s* (sum), HPE estimate using the `hpe()` function in the `fasthplus` R package, and HPB estimate using the `hpb()` function in the `fasthplus` R package for increasing sizes of datasets with *n* = 100, 500, 1,000, and 3,000 observations and 500 features. All observations were simulated from *N*_500_(0, 1) and then split evenly in two groups. The hpb procedure used *r* = 0.05 × *n* with *t* = 30, and the `hpe` procedure used *p* = 1001 with the grid search algorithm.

| obs | 100 | 500 | 1000 | 3000 |
| --- | --- | --- | --- | --- |
| dis | 0.01 | 0.22 | 3.00 | 36.71 |
| adj | 0.00 | 0.01 | 0.08 | 0.44 |
| sum | 0.08 | 59.68 | NA | NA |
| hpe | 0.12 | 0.15 | 0.28 | 1.86 |
| hpb | 0.01 | 0.05 | 0.20 | 5.74 |

